# MAGScoT - a fast, lightweight, and accurate bin-refinement software

**DOI:** 10.1101/2022.05.17.492251

**Authors:** Malte Christoph Rühlemann, Eike Matthias Wacker, David Ellinghaus, Andre Franke

**Affiliations:** Institute of Clinical Molecular Biology, Kiel University, Kiel, Germany; Institute for Medical Microbiology and Hospital Epidemiology, Hannover Medical School, Hannover, Germany

## Abstract

We introduce `MAGScoT`, a fast, lightweight and accurate implementation for the reconstruction of highest-quality metagenome-assembled genomes (MAGs) from the output of multiple genome-binning softwares. `MAGScoT` outperforms popular bin-refinement solutions in terms of quality and quantity of MAGs as well as computation time and resource consumption. `MAGScoT` is available via GitHub (https://github.com/ikmb/MAGScoT) and as an easy-to-use Docker container (https://hub.docker.com/repository/docker/ikmb/magscot).

## Main text

Reconstructing high-quality microbial genome sequences from complex metagenomic sequencing libraries is crucial for studying microbial traits and their influence on host health or disease, or their role in adaptation to changing environmental factors. A typical workflow for this task usually encompasses the assembly of larger contiguous sequences from short-read sequencing libraries, followed by binning of these contigs into larger sets called metagenome-assembled genomes (MAGs), and has been the basis for large-scale metagenomic surveys. ^1,2^

Binning of contigs into MAGs can be achieved by one or more specific properties of the assembled contigs, i.e. similarity in coverage, k-mer or tetranucleotide frequencies, GC content or the presence of specific marker genes, paired with different types of statistical and/or machine learning approaches for clustering contigs into bins, such as expectation maximization or variational autoencoders.^3–6^ The resulting MAGs are subsequently assessed for their quality, e.g. by the `CheckM` software,^7^ which uses clade-specific microbial marker genes to assess the completeness of a MAG, as well as contamination due to incorrect assembly or binning, and the presence of marker gene duplicates that may indicate the binning of multiple closely related genomes.

It was demonstrated that applying and combining multiple binning approaches can be useful to reconstruct more and better quality MAGs from metagenomic datasets.^8^ Two popular algorithms for such a bin-refinement are `DASTool` and the bin-refinement module of the `metaWRAP` pipeline.^8,9^ Briefly, `DASTool` relies on an iterative dereplication, aggregation and scoring algorithm that selects the highest-quality genome from a collection of contig-to-bin mappings based on 51 bacterial- and 38 archaeal-specific single-copy marker genes. The `metaWRAP` refinement module, henceforth simply called `metaWRAP,` relies on the identification of overlaps between multiple bin sets, from which it constructs potentially higher quality hybrid bins. The `CheckM` software is then used to select the highest quality version of a genome from original and hybrid bins. `metaWRAP` can use the output of up to three individual binning softwares at once; if more input sets are to be included, they must be distributed across multiple `metaWRAP` runs.

We introduce here the tool `MAGScoT` (= MAG Scoring Tool) which combines these two concepts into a single approach. `MAGScoT` relies on two sets of microbial single-copy marker genes from the Genome Taxonomy Database Toolkit,^10^ 120 bacterial and 53 archaeal, stored as HMM-profiles for fast annotation of amino acid sequences predicted from the assembled contigs.^11–13^ Presence profiles of marker genes are compared between the results of the individual binning algorithms, and new hybrid candidate bins are created if MAGs from binsets share a user-adjustable proportion of marker genes (default: 80%). Subsequently, all original bins from each binning software, and the newly constructed hybrid bins are scored in the same way as introduced by `DASTool`, with the option to weight completeness (default: *a*=1) and penalize contamination (default: *b*=0.5 and *c*=0.5). `MAGScoT` is not limited in the number of binning algorithms outputs that can be used as input. Details on the `MAGScoT` algorithm can be found in the **Online Methods** section.

We evaluated the binning performance and computational demand of `MAGScoT`, `DASTool` and `metaWRAP` on two datasets. The first dataset was the simulated “marine” dataset of the CAMI2 challenge (see **Methods**).^14^ This complex dataset with known ground truth offered a comparison to a perfect binning outcome. The second dataset was a dataset of 50 randomly selected metagenome samples from the integrative Human Microbiome Project (iHMP/HMP2, see **Methods**), which allowed us to compare performance in a real-world setting.^15^ All three bin-refinement softwares were run on the same input data derived from four binning algorithms: `MaxBin2`, `MetaBAT2`, `CONCOCT` and `VAMB` (**Supplementary Tables 1 & 2**). All original and refined bins were evaluated using `CheckM`.

Computational time and hardware demand were assessed on a single compute node in a high-performance computing environment in individual jobs restricted to 8 CPU cores and 80 GB of RAM (**Table 1, Figure 1, Supplementary Tables 3 & 4**). The computation time was longest for `metaWRAP` in both datasets due to the iterative usage of the `CheckM` software to score the individual bins. Of note, RAM usage was also highest for `metaWRAP,` as `CheckM` relies on large reference trees for scoring based on clade-specific marker genes. `MAGScoT` showed the fastest performance of all tools in both datasets. Especially in the highly complex and fragmented synthetic “marine” dataset, HMM-based annotation and the main `MAGScoT` algorithm were approx. 15-fold faster than `DASTool` for the equivalent steps, with both programs using the same predicted amino acid sequences as input (**Table 1**). The run time for gene/protein prediction from contigs and HMM-based annotation for marker genes scales linearly with metagenome complexity; however, additional compute resources can speed-up these steps almost linearly.

**Table 1:**
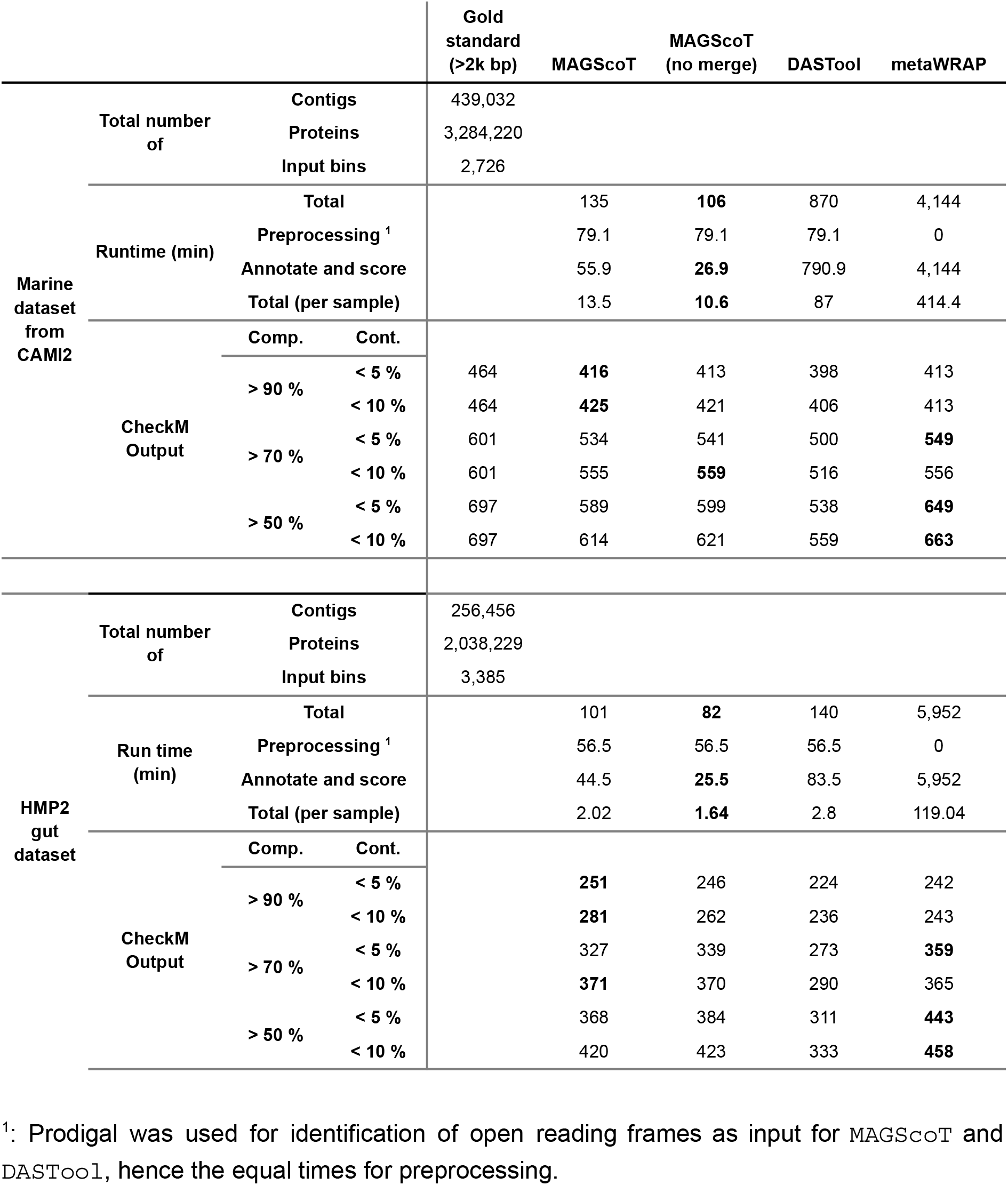
Overview of bin-refinement performances in the marine and HMP2 gut metagenome datasets. Performance was evaluated by runtime on a single compute node in a high-performance computation enviroment with 8 CPUs and 80Gb of RAM and the quality and quantity of recovered MAGs based on CheckM scores for clade-specific completeness (Compl.) and contamination (Cont.).

**Figure 1:**
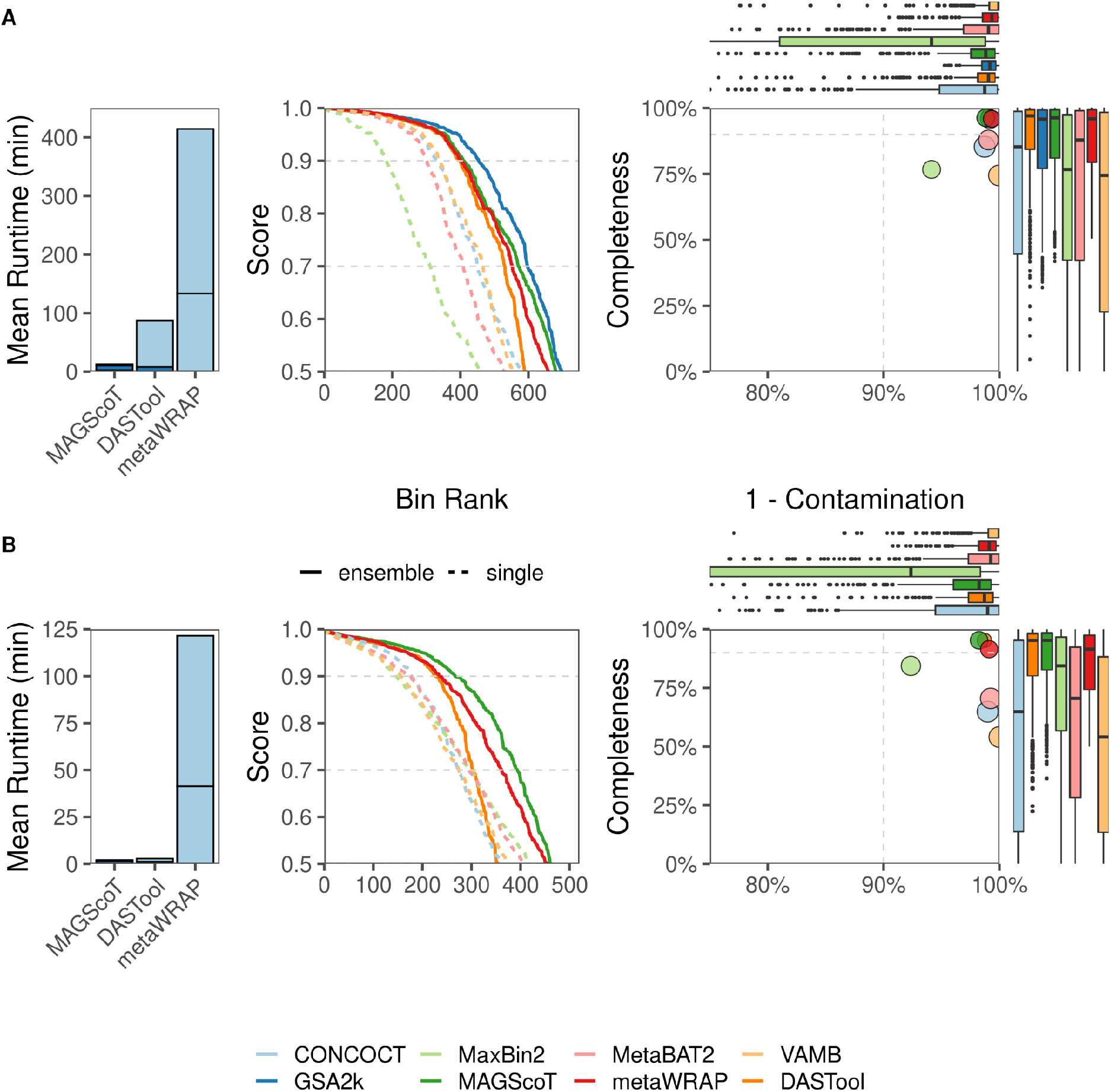
Summary of binning performance in the (A) simulated marine dataset from the CAMI2 challenge and the (B) collection of 50 gut metagenomes from the HMP2 based on CheckM output. The left panels show mean runtime per sample on 8 CPUs and 80 GB of RAM. In the center panel, completeness and contamination are combined into a score (= completeness - 0.5 × contamination), which was used to rank the bins. `GSA2k` (dark blue) in the simulated marine dataset represents the perfect-case scenario that can be obtained from the gold standard assembly filtered by a minimum contig length of 2000bp. The right panel summarizes the median completeness and contamination by binning or refinement software. Boxplots depict the distribution of binning completeness and contamination with each software output. The dashed gray lines in the center panels show proxies for high quality (score>0.9) and good (score>0.7) MAGs. The dashed gray lines in right panels show the completeness and contamination thresholds for high quality MAGs at 90% and 10%, respectively.

For all three bin refinement algorithms, the completeness and contamination statistics showed very good values, with the respective median values clearly surpassing the thresholds for HQ MAGs and showing performance close to the gold standard for the simulated marine dataset (`GSA2k`, **Figure 1**, right panels). The statistics clearly show the importance of refining the bins using the results of multiple binning algorithms to leverage their respective strengths, i.e. `MaxBin2` showed the highest completeness scores of all individual binners in the real metagenomic data, but at the expense of the highest contamination. Conversely, `VAMB` reconstructs bins with very low contamination at the cost of the lowest overall completeness. All `CheckM` scores for all original and refined bins can be found in Supplementary **Tables 5 and 6**.

In both the synthetic marine and the real-world gut metagenome datasets, `MAGScoT` produced the highest number of high quality MAGs (HQ, completeness > 90%, contamination < 10%, **Table 1, Figure 1**, middle panels). The number of medium quality (MQ) MAGs (completeness > 50 %, contamination < 10%) was highest for `metaWRAP`. It is likely that `MAGScoT` identifies fewer MQ MAGs since its default scoring cutoff is 0.5, which is calculated based on bin completeness and contamination (see **Methods**), thus bins marginally above a completeness of 50%, even with low contamination up to 10%, are not reported as they fall below this threshold. Comparing the `MAGScoT` output to the gold standard for the simulated marine community, `MAGScoT` identified 425 of 464 (92.6%) possible HQ MAGs and 614 of 697 (88.1%) of MQ MAGs (**Table 1**). Disabling the bin merging steps in `MAGScoT` resulted in the lowest computation time of all bin-refinement software, accompanied by a slight decrease in the number of HQ MAGs but an increase in MQ MAGs (**Table 1**).

In summary, `MAGScoT` outperforms the two popular bin refinement solutions `metaWRAP` and `DASTool` in terms of the quantity and quality of MAGs recovered, while being more economical with computational resources.

## Data availability

All data were required from publicly accessible repositories (marine synthetic community data: https://frl.publisso.de/data/frl:6425521/marine/short_read/; iHMP/HMP2 data: https://ibdmdb.org/tunnel/public/HMP2/WGS/1818/rawfiles).

## Code availability

MAGScoT is available via GitHub (https://github.com/ikmb/MAGScoT). All scripts to produce the binning results and the subsequent refinement are also available via GitHub (https://github.com/mruehlemann/MAGScoT_benchmark_scripts).

## Online Methods

All scripts used for processing of the data are available via Github: https://github.com/mruehlemann/MAGScoT_benchmark_scripts

### Dataset acquisition and processing

Short reads and gold standard assembly contig data of the simulated marine dataset was downloaded from the CAMI ressource server (https://frl.publisso.de/data/frl:6425521/marine/short_read/). Gut metagenome short read data for the HMP2 gut dataset were acquired from the servers of the Inflammatory Bowel Disease Multi’omics Database (https://ibdmdb.org/tunnel/public/HMP2/WGS/1818/rawfiles). Here, data of the first 100 samples were downloaded and the 50 samples with the most reads were kept to ensure high enough read depth for the subsequent steps. Short read data was assembled using the Megahit software to acquire metagenomic contigs. From here, both datasets were treated equally. Contig data was filtered to retain contigs with a size of >= 2000 bp, as this was used as minimum contig size threshold for all binning softwares. For each sample, reads were mapped against the “own” contigs using minimap2 (version 2.17-r941). In addition, for both datasets, a cross-sample contig catalog was created and also the reads of the respective datasets were mapped against this catalog using minimap2. The resulting mapping files (SAM format) were sorted and converted to binary (BAM) format using Samtools (version 1.9). The jgi_summarize_bam_contig_depths executable from the MetaBAT2 binning algorithm was used with default options to receive coverage information for the individual sample mappings and the mappings to the dataset catalogs. Per-sample contig sequences and mapping results were used for the MetaBAT2, Maxbin2 and CONCOCT binning softwares with default options. The cross-mappings to the dataset catalogs were used in VAMB with default options and using GPU-accelerated calculations. The binning outputs were used in the bin-refinement algorithms.

As ground truth data was available for the simulated marine dataset, best-case binning results were derived from these profiles. To have a fair comparison of binning tools and refiners to the ground truth with a focus on bacterial and archaeal genomes, the original metagenomic contigs were filtered to include only data from bacterial/archaeal genomes which were represented to at least 40% in contigs of >= 2000 bp, as this was the length cutoff used for the binning algorithms. In addition, only contigs >= 2000 bp were included in the ground truth bin quality assessment using CheckM. The filtered gold standard assembly (gsa2k)

### Binning algorithms

This section gives a brief summary of the four used as baseline for the bin refinement:

**MetaBAT2** (v2.12.1): MetaBAT2 relies on normalized tetranucleotide frequencies (TNFs) calculated from the input contigs to create an initial graph for graph-based clustering. Subsequently, similarity scores from TNFs and correlation between contig abundances are used iteratively to be incorporated into the graph structure. A label propagation algorithm is used to score connections and identify clusters of contigs.

**MaxBin2** (v.2.2.7): In addition to TNFs and contig abundances MaxBin2 uses a set universal single-copy marker genes for the identification and clustering of metagenomic bins. The clustering is based on an Expectation-Maximization algorithm.

**CONCOCT** (v.1.1.0): CONCOCT more generally uses k-mer frequencies and contig abundances for clustering, by default, the k-mer length in CONCOCT is 4, which makes it congruent to the aforementioned TNFs. One major difference, is CONCOCT suggests to cut contigs into pieces of 10kb, to identify and disentangle mis-assembles. The clustering of the contigs into bins is done by a Gaussian mixture model fit with a variational Bayesian approximation.

**VAMB** (v.3.0.3): VAMB is the most recent of the herein used binning algorithms. Similar to the other binning tools, VAMB relies on contig coverage and TNFs. The major difference to the other binning tools is the use of unsupervised deep learning via variational autoencoders and can be accelerated by the use of GPUs.

### MAGScoT algorithm

The MAGScoT algorithm is implemented in R (v.4.0) and by default takes two input files: 1. the HMMER-based GTDBtk bacterial and archaeal single-copy marker gene (SCG) annotations (release 207) of amino acid sequences of open reading frames from the assembled contigs identified by prodigal (option: -p meta) and 2. the contig-to-bin mappings of the individual binning algorithms.

In a first step, MAGScoT pairs annotated marker genes with their respective contigs. Subsequently, bins with a predefined minimum number of identified marker genes (default: 20) are scanned for overlap with all other bins based on shared genes which were annotated as marker genes. If this overlap surpasses a threshold (default: 80%), a hybrid bin of the two original bins is added to the total bin collection. This process can be performed iteratively in which also the newly formed hybrid bins are scanned for overlap with additional original bins. By default, this is done twice, as each iteration can add hundreds of new bins for the scoring and refinement process. The creation of hybrid bins can be skipped by the --skip_merge_bins flag when starting MAGScoT.

The second step is the scoring and refinement iteration. Here, each bin is scored using sets of SCGs, 53 for archaea and 120 for bacteria. The scoring function is the same as in DASTool:

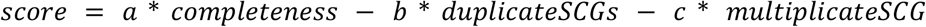

with:

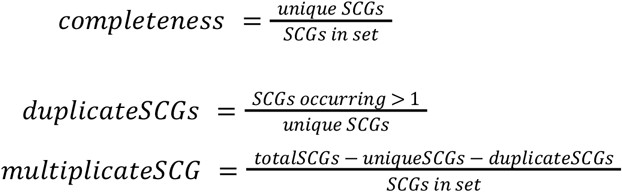

The scoring weights are *a*=1, *b*=0.5, *c*=0.5 by default, but can be adjusted by the user, e.g. to emphasize completeness. The higher score from the archaeal and bacterial sets of SCGs is used as the bins score. Each round, the bin with the highest score is picked from the dataset. A major reason for the speed of MAGScoT is that only remaining bins that changed due to the removal of the winning bin are re-scored. Additionally, after each scoring, the reporting threshold *t* (default: 0.5) is used to remove bins from the dataset based on if their completeness does not surpass *t*, as bins below this completeness will never reach the reporting threshold. The scoring and refinement process is repeated until there is no more bin surpassing *t*.

### Refinement benchmark

The same input generated by the individual binners was used as input for MAGScoT, DASTool and metaWRAP. DASTool was run with default options, however using the -p option to use the same amino acid sequences as used in MAGScoT and to have a better estimate of the actual time DASTool spends in the iteration and scoring phase. As the bin-refinement module of metaWRAP accepts a maximum of three bin sets simultaneously as input, it was first run with output from MaxBin2 and VAMB. In a second run, the resulting refined bins were used as input together with the output from CONCOCT and MetaBAT2. This strategy is recommended by the metaWRAP developers (see https://github.com/bxlab/metaWRAP/blob/master/Usage_tutorial.md#step-5-consolidate-bin-sets-with-the-bin_refinement-module). After refinement, all original and refined bins were scored using CheckM. The CheckM scores for all bins can be found in Supplementary Tables 5 and 6 for the simulated marine and the real HMP2 gut metagenomes, respectively. Running times for all samples and all steps in the pre-processing and main bin-refinements softwares can be found in Supplementary Tables 3 and 4.

